# Swimming with the current improves juvenile survival in southern elephant seals

**DOI:** 10.1101/2023.11.05.565668

**Authors:** Dahlia Foo, Clive R. McMahon, Mark A. Hindell, Michael Fedak, Martin Biuw, Bernie McConnell, Ben Raymond

## Abstract

Understanding juvenile survival is crucial for the population ecology of long-lived species, where parental guidance can significantly influence survival rates of completely naive juveniles. In southern elephant seals (*Mirounga leonina*), however, offspring receive no knowledge from their parents, only fat reserves. This research focuses on how dispersal direction on their maiden foraging trip and physical traits influence the survival of naïve southern elephant seal pups at Macquarie Island. We tracked 44 pups with satellite tags during their post-weaning migration and compared their movements to the post-moult winter migrations of 58 adult females. We found that most pups (61.2%) travelled southeast, in line with the Antarctic Circumpolar Current. Pups travelling with the predominant east-southeast current had a 1.5 times higher survival rate for their first trip than those swimming westward against it. Those that swam with the current and were heavier were more likely to survive their first year. Adult females showed different dispersal patterns, where they travelled southwards towards Antarctic waters, implying that learning from experience influences their direction. Future investigations into the role of the primary eastward current in the sub-Antarctic on circumpolar movement patterns of top marine predators could expand our understanding of Southern Ocean ecology.

## Introduction

Explaining why young animals die is crucial for both studying how populations change over time and the survival of long-lived and slow-reproducing species [1,2]. One of the key factors influencing juvenile survival is their ability to forage effectively [3,4]. Young animals, especially in their first year, face numerous challenges in securing food, often due to their physical limitations and inexperience [5,6]. This vulnerability is evident in the high mortality rates observed among both juvenile birds [3,7] and mammals [6,8].

In numerous mammal species, survival often hinges on maternal guidance [9,10], with a range in terms of transfer of knowledge from adult to offspring. On one end, in species such as elephants [11], chimpanzees [12], and dolphins [13], social development and the transfer of knowledge from parents to offspring is particularly important. Somewhere in the middle, in seal species such as harbor [14], bearded [15] and Weddell seals [16], there is some maternal supervision of their offspring at-sea where pups may indirectly benefit by observing foraging locations and preferable prey. At the other extreme, juveniles may become independent without maternal knowledge transfer, relying instead on inherited behavioural traits, or experienced gained in early life.

In the case of southern elephant seals (*Mirounga leonina*), apart from maternal investment through lactation (during which pups never enter the water), there is no further guidance or knowledge transfer from the mothers. Once mothers leave their pups at 24 days of age, their pups are completely naïve concerning foraging behaviour and areas and are left to fend for themselves [17]. Weaning mass is therefore a crucial factor for first-year survival in southern elephant seals, influenced significantly by the mother’s foraging success during gestation [1,18,19]. Heavier weaned pups tend to have higher survival rates [20], likely because they start off with more stored energy from their fat reserves [21]. This extra energy acts like a safety net, giving them more time to find their first meal and learn how to forage effectively [22,23].

Adults and juvenile elephant seals tend to forage in different areas [24,25], raising questions about the developmental shifts in innate and learned behaviours over time. However, the mechanisms that guide these inexperienced pups to their feeding grounds are not well-understood. Current theories suggest a mix of innate navigational capabilities and learned foraging behaviours [26,27]. This makes elephant seal pups an excellent model for investigating the balance between innate traits and learned skills in determining an individual’s fitness, especially when there is no direct transfer of informational knowledge from parent to offspring.

Southern elephant seals breed on sub-Antarctic islands and have a circumpolar distribution [28]. They forage in a variety of regions within the Southern Ocean, particularly in inter-frontal zones where their prey – mesopelagic fish, squid, and crustaceans like krill [29–32] – tends to be sparsely and unevenly distributed [33]. These zones, made up of different water masses, are influenced by the Antarctic Circumpolar Current (ACC), which is the dominant eastward-flowing current in the Southern Ocean ecosystem [34,35].

Optimal foraging theory suggests that animals should adopt foraging strategies that maximise their net energy intake – this means maximising energy gains and minimising energy costs [36]. In the case of naïve southern elephant seal pups, without knowledge of where prey is located, we hypothesise that their movement within currents might not be a deliberate strategy to conserve energy. Instead, due to their random directional movement in a current, they end up being transported further downstream. This passive transport may reduce the energetic cost of travel during their vulnerable early weeks and potentially optimise foraging opportunities [37–39]. Because seals forage in a patchy environment, they often need to travel relatively long distances to find food [35]. Thus, whether intentional or not, being carried by the currents can be beneficial, especially in the early stages of a juvenile’s first trip to sea when they are losing body condition [23].

In this study, we used a unique dataset of individually marked weaned elephant seal pups tracked during their first foraging trip and their subsequent survival to 1) determine where the seals go on their first foraging trip and if they followed oceanic currents, 2) compare where they go to adult females, and 3) relate their movement to their first trip and first year survival. The first aim informs our understanding of how innate abilities and environmental cues guide naïve pups’ foraging patterns. The second aim provides insights into the developmental process and influence of innate versus learned behaviours. Finally, the third aim will correlate movement patterns and survival outcomes, providing a comprehensive view of the challenges and strategies in the early life stages.

## Methods

### Data collection

Sixty-nine weaned pups at Macquarie Island (54°30′ S, 158°57′ E) were tagged with ARGOS satellite relayed data loggers (SRDL, Sea Mammal Research Unit) during their post-weaning fast in December 1995, 1996, 1999 and 2000. These SRDLs provided locations and dive metrics. The pups were also flipper tagged (Jumbo Rototag, Dalton Supplies, Henley-on-Thames, UK) and permanently marked by hot iron branding for future identification as part of a long-term demographic study [2,40,41]. Researchers made resights of marked animals every year at Macquarie Island the resight dataset spanned from 23 November 1951 – 1 October 2014. Details on how seals were captured, weighed, and handled can be found in Hindell et al. [17].

It is important to note that the weaners in our dataset were not selected randomly. Instead, the aim was get a range of weaner sizes [18]. This may lead to some bias in our results, however given our relatively large sample size and the biological context of this research, we believe that our results are robust.

We also obtained 67 adult female post-moult winter migration tracks from Macquarie Island which were collected from February–October in 2000–2005 and 2010 [42]. Out of the 67 tracks, 30 were obtained by SRDLs and the remaining 37 by geolocation light loggers (GLS; from time-depth recorders, Wildlife Computers, Redmond, USA). GLS tags are accurate to ∼70 km [43]. Some of these tracks did not start at the colony and were removed from further analyses.

### Processing tracks

Only seals that left the colony and provided > 10 days of ARGOS data were kept for further analysis. Additionally, some seals had missing wean mass in the dataset (n = 3) – these were also removed from further analyses. The at-sea locations provided by Argos were filtered using a correlated random walk state-space model with a 4 m s^-1^ max velocity threshold via the R package *foieGras* [44] and its successor *aniMotum* [45]. We used a 12 h time step and to provide two location estimates per day to simplify the data set while still capturing the essential movement patterns of each seal.

### Identifying foraging trips

At-sea locations within 50 km of the colony were considered to be at the colony [21] as seals may spend time in the water close to the colony to thermoregulate rather than forage. Foraging trips were determined as trips that lasted > 1 day. Complete foraging trips were determined as trips that started and ended at the colony. Only the first long foraging trip (>30 days) of individuals were used for further analysis.

### First foraging trip and first year survival

A seal was considered to have survived its first foraging trip if there was either a complete track of this trip or if the seal was seen again at the colony during the resight period. The latter would indicate that the tag failed before it returned to the colony during its first foraging trip. Additionally, seals were only considered to have survived their first year of life if they were resighted at least once after 1 year of being tagged until the end of the study in 2014.

### Determining dispersal bearing

To determine the direction that the seals travelled when they left the colony, we calculated the bearing between Macquarie Island Isthmus and the last location on day 5 of their foraging trip. Day 5 was chosen as by this time seals would have travelled sufficiently far away from the colony.

### Particle trace

To investigate if weaners were following the flow of the current when they left the colony, we simulated the path that a particle would take if it were dropped at the starting location on the same day that each seal left for their first foraging trip. We used the *currently* R package [46] to generate a particle trace (“track”) with a 12-h timestep, for each weaner based on relevant parameters (starting location, duration, departure date) of their first foraging trip.

To ensure a robust and accurate comparison between seal and particle dispersal, we used the particle’s location closest to the corresponding seal’s position on the fifth day of its trip as the reference point for calculating the dispersal bearing from Macquarie Island. This approach was chosen because a preliminary visual inspection revealed that particle traces could diverge from their initial direction over time, making the most distant point a less reliable metric for comparison (see Supplementary Figure 1). The predominant flow of the ocean current was towards the east-southeast. Hence, seals which dispersed in that direction were classified as travelling with the current.

### Statistical analysis

Angular/directional data were analysed using a circular statistics R package, *circular* [47]. To test whether mean bearings differed between groups of seals, we used Watson’s two-sample test of homogeneity [48]. To test whether the seals’ bearings were uniformly distributed, or whether a preferred direction of travel was evident within a certain group, we used the Rayleigh test of uniformity.

Logistic regression models for binomial data were fitted to test the likelihood of first trip and first year survival as response variables against year and the interaction term between wean mass and if seals swam with the flow of the currents. We did not include sex because it has little effect on first year survival [41] and including it would have reduced the power of our dataset. Using the *MuMIn* R package [49], candidate models were ranked by AICc, and those with delta AICc < 2 were averaged to generate the final model. Model assumptions were checked by simulating residuals of the global model 1000 times using the *DHARMa* R package [50]. All results are reported as means ± standard error unless otherwise stated.

## Results

### Overview of tracking and survival data

We used data from 44 weaners and 58 adult females in the final analyses. The mean birth date of weaners was 1 October ± 1 d. Their mean weaning date was 1 November ± 2 d (Table 1). The median departure days of the year for weaners and adult females were December 18 ± 11 d and February 10 ± 8 d, respectively. The overall mean duration of the first foraging trip of weaners was 113 ± 11.6 d (range = 6–340 d) and their mean maximum distance travelled from the colony was 1807 km ± 134 (range = 231–3931 km). Twenty eight out of 44 (64%) weaners survived their first trip. Most (80%) weaners travelled southeast from Macquarie Island during their first foraging trip, while few (13%) travelled southwest (Figure 1). In comparison, adult female tracks had a more even and broader southern distribution (Figure 1).

**Figure 1.**
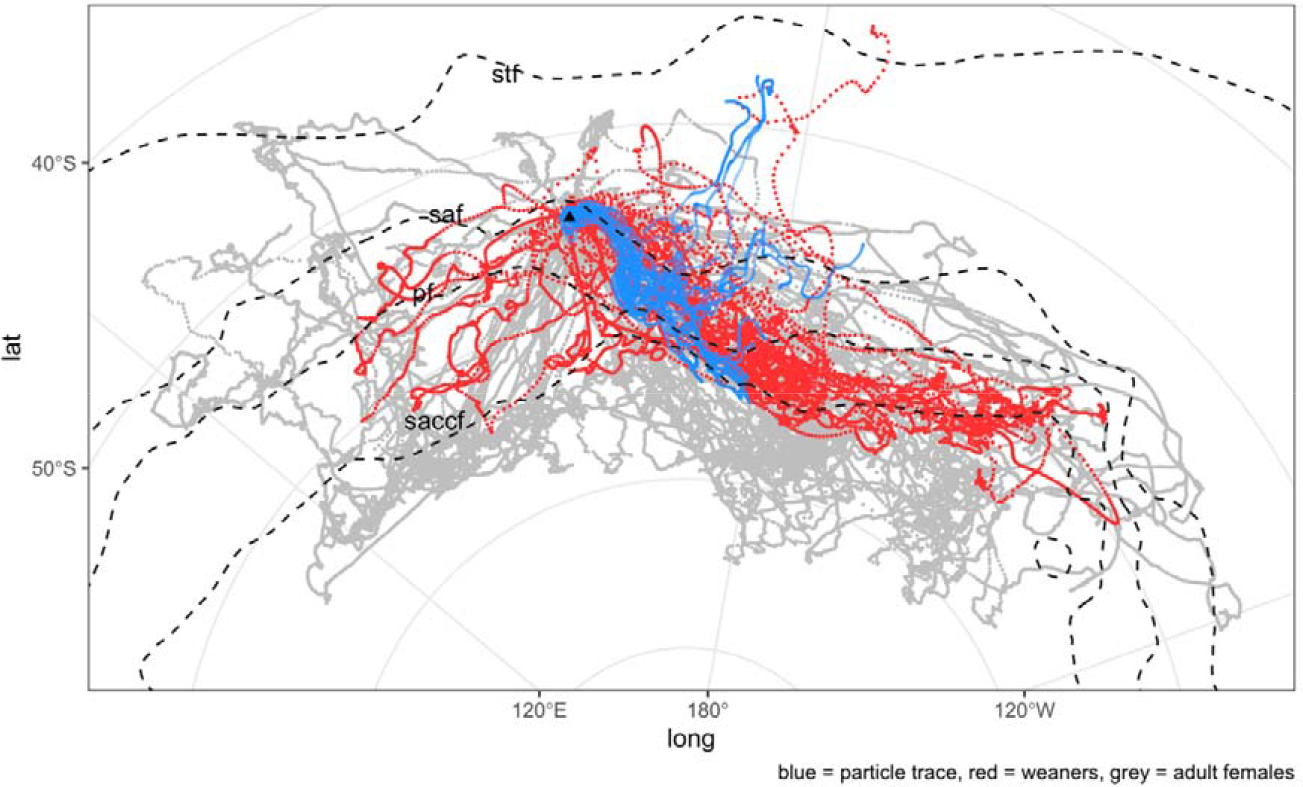
Tracks of particles (blue), weaned elephant seals (red), and adult female elephant seals (grey) from Macquarie Island (triangle). Also shown are the four major oceanic fronts in the region: subtropical front, stf; sub-Antarctic front, saf; polar front, pf; southern Antarctic circumpolar current front, saccf.

**Table 1.**
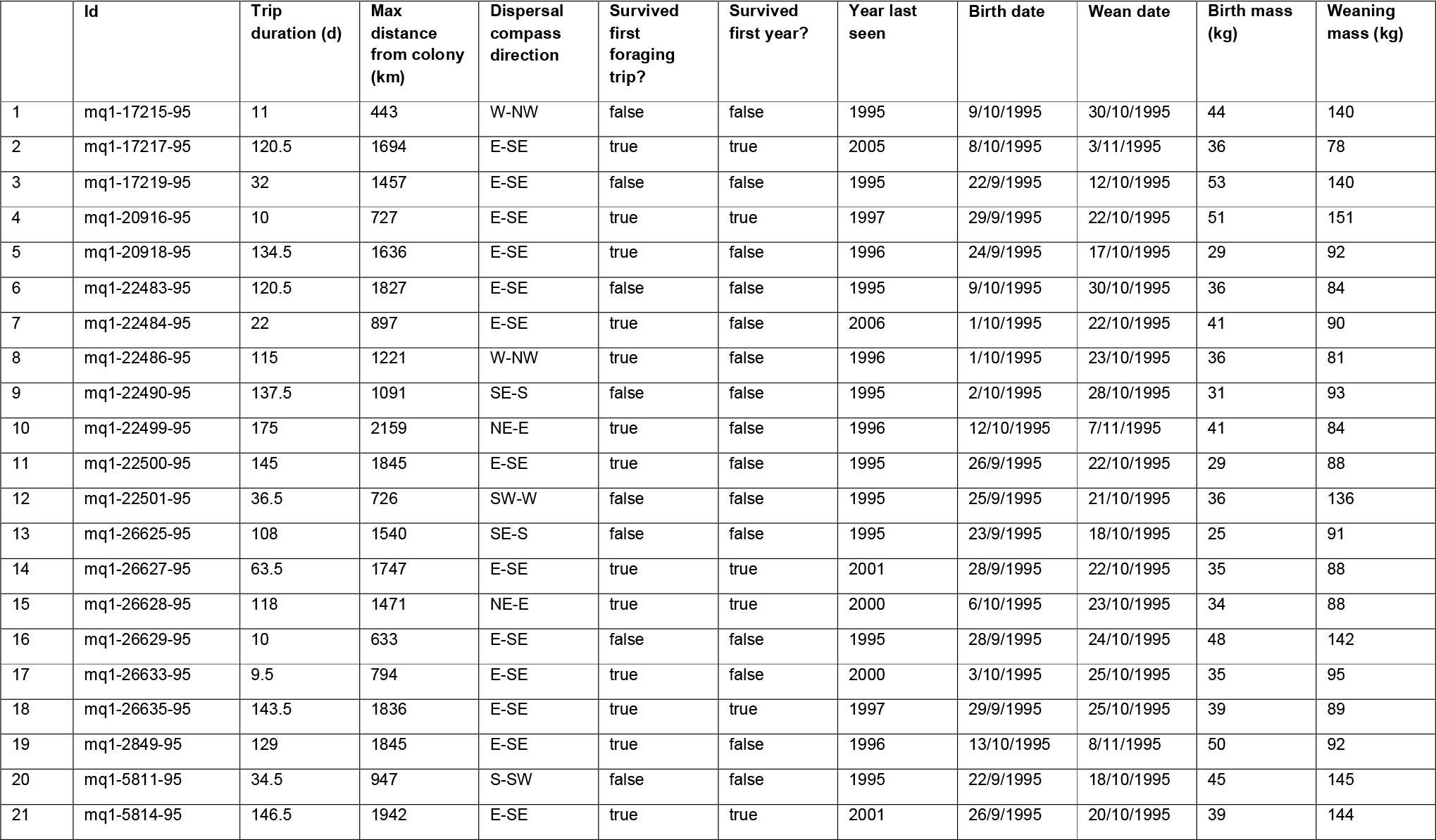
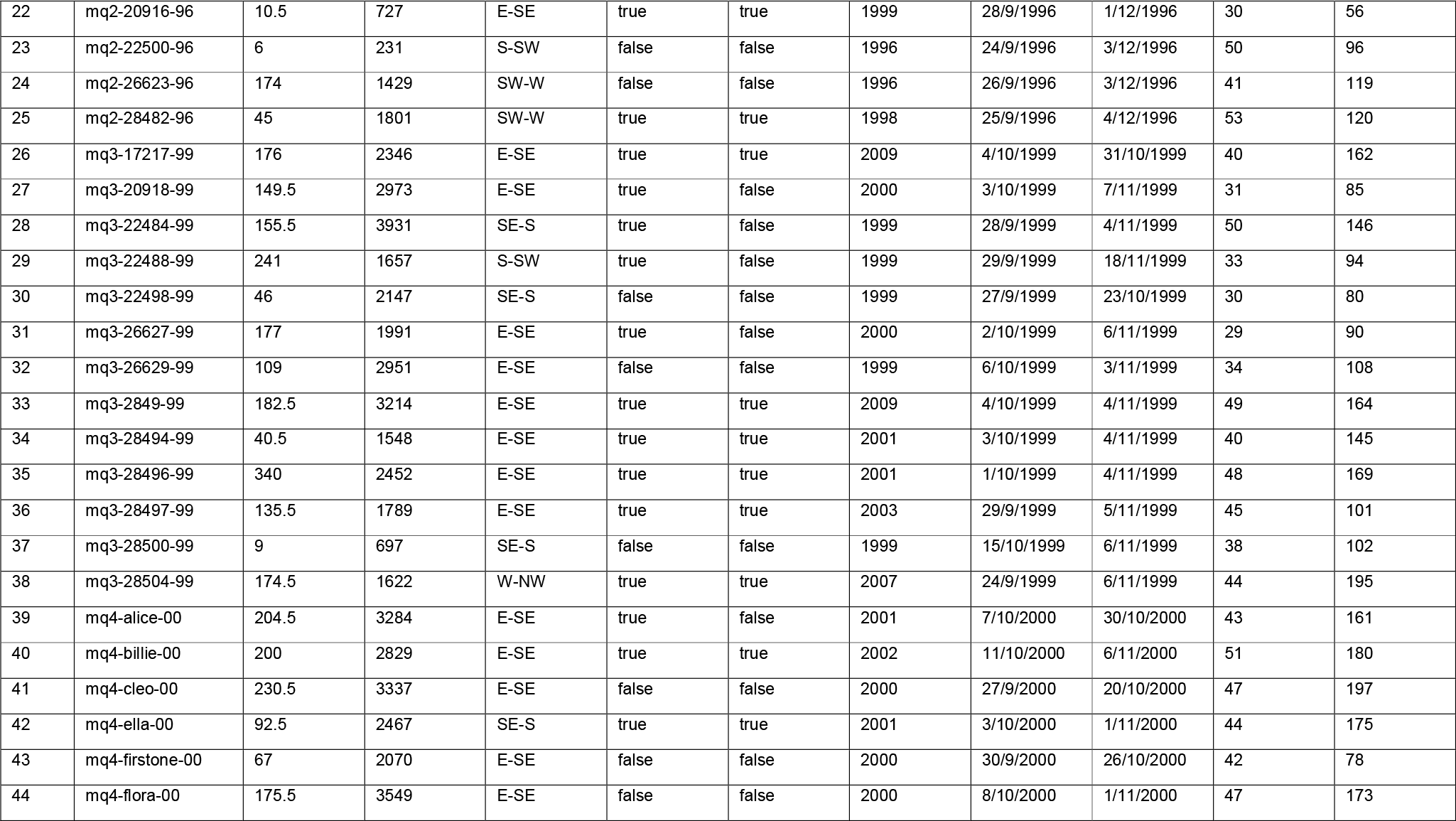
Summary of the first foraging trip deployment and life history information of individual weaners.

### Weaners and adult females dispersal directions

The overall mean distance from the colony of the reference location (i.e. 5 days after departure) used to calculate dispersal bearing for weaners, particle traces, and adult females were 368 ± 19 km, 313 ± 26, and 422 ± 17 km, respectively (Supplementary Table 1). Adult females and weaners dispersed from the colony in different directions (Figure 1 and Figure 2; adult female bearing = 167.1 ± 0.1°; weaner bearing = 126.5 ± 0.1°; Watson’s two sample test: test statistic = 1.13, p < 0.001; Supplementary Table 1). There was no difference between the mean weaner and particle trace bearings (Figure 2; Supplementary Table 1; particle trace bearing = 116.8 ± 0.1°; Watson’s two sample test: test statistic = 0.07, p value > 0.10) and between adult female and particle trace bearings (Watson’s two sample test: test statistic = 1.31, p-value < 0.001). Both adult females (Rayleigh test of uniformity: test statistic = 0.713, p = 0) and weaners (Rayleigh test of uniformity: test statistic = 0.623, p = 0) had preferred dispersal bearings i.e. the bearing was uniform among individuals in the group.

**Figure 2.**
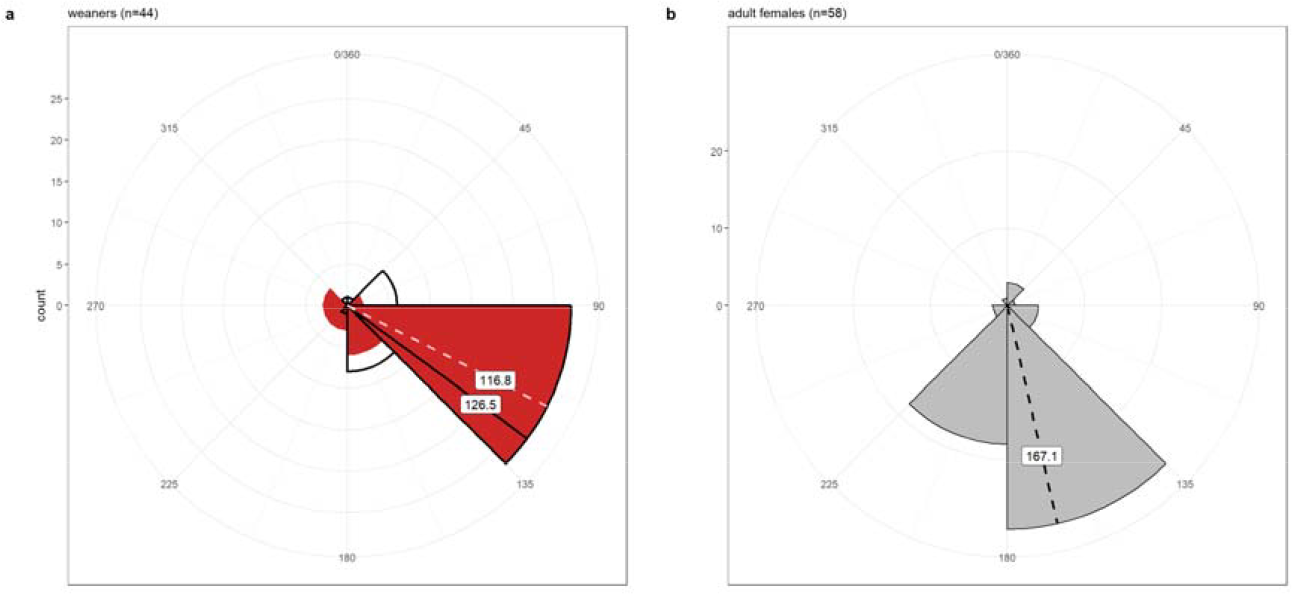
Circular histograms of the dispersal direction of (a) weaners in red and their corresponding particle trace in black outline, and (b) adult females. Dashed (weaner and adult female) and solid (particle trace) lines show the mean bearing which is labelled.

### Effect of wean mass and dispersal direction on survival

Out of 44 weaners, 27 (61.2%) swam east-southeast (with the current), of which 20 survived (74.1% survival rate). Of the remaining 17 weaners that swam against the current, only 8 survived (47.1% survival rate). Moreover, 12 weaners (44.4% survival rate) that swam with the current in their first trip survived their first year, while only 4 weaners (23.5% survival rate) that did not swim with the current in their first trip survived until their first year.

Weaners were more likely to survive their first foraging trip if they dispersed from the colony with the flow of the predominant current (east-southeast direction) (Table 2; Figure 3; Figure 4; Supplementary Table 2). Similarly, they were more likely to survive their first year of life if they dispersed with the flow of the predominant current during their first foraging trip and had a heavier weaning mass (Table 2; Supplementary Table 3). For weaners that did not disperse with the flow of the current, the effect of wean mass on first year survival was greater i.e. steeper slope of model fit (Figure 5). Additionally, there was no difference in weaning mass between seals that went with the flow versus those that did not (ANOVA test: F_1,42_ = 0.08, p = 0.7).

**Table 2.** Summary of the first trip survival and first year survival logistic generalised linear models.

|  | Estimate | SE | Adjusted SE | z-value | p-value |
| --- | --- | --- | --- | --- | --- |
| **1st trip survival** |  |  |  |  |  |
| (Intercept) | 0.13 | 0.54 | 0.55 | 0.24 | 0.81 |
| is_ESE (true) | 1.17 | 0.65 | 0.67 | 1.73 | 0.08 |
| **1st year survival** |  |  |  |  |  |
| (Intercept) | -2.35 | 1.95 | 1.98 | 1.19 | 0.23 |
| weanmass | 0.02 | 0.01 | 0.01 | 1.30 | 0.19 |
| is_ESE (true) | 1.72 | 2.07 | 2.11 | 0.81 | 0.42 |
| weanmass:is_ESE (true) | -0.02 | 0.02 | 0.02 | 1.06 | 0.29 |

**Figure 3.**
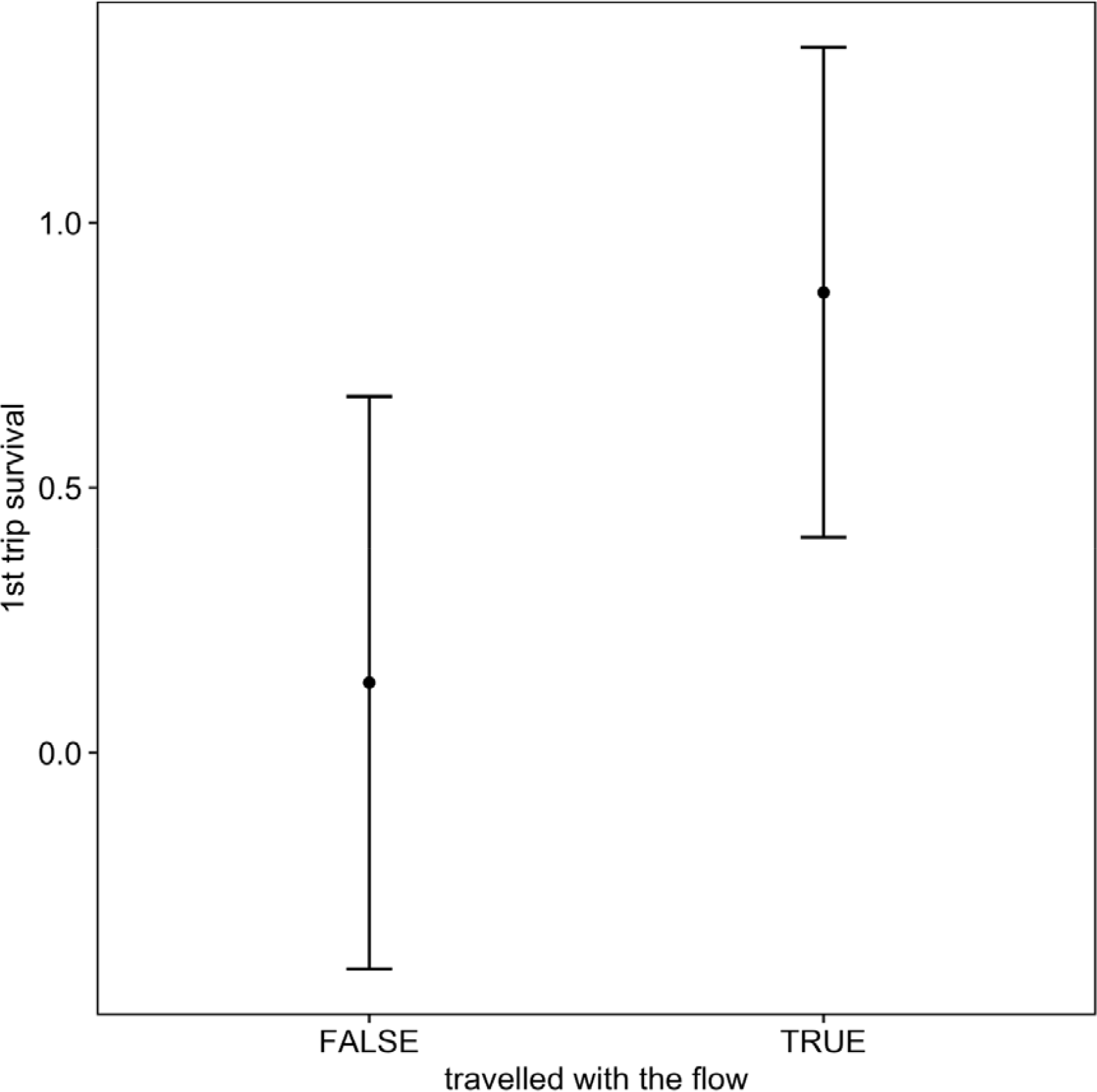
Predicted effects of whether southern elephant seal weaners dispersed in the east-southeast sector direction on first trip survival. Solid circle = mean; error bars = SE.

**Figure 4.**
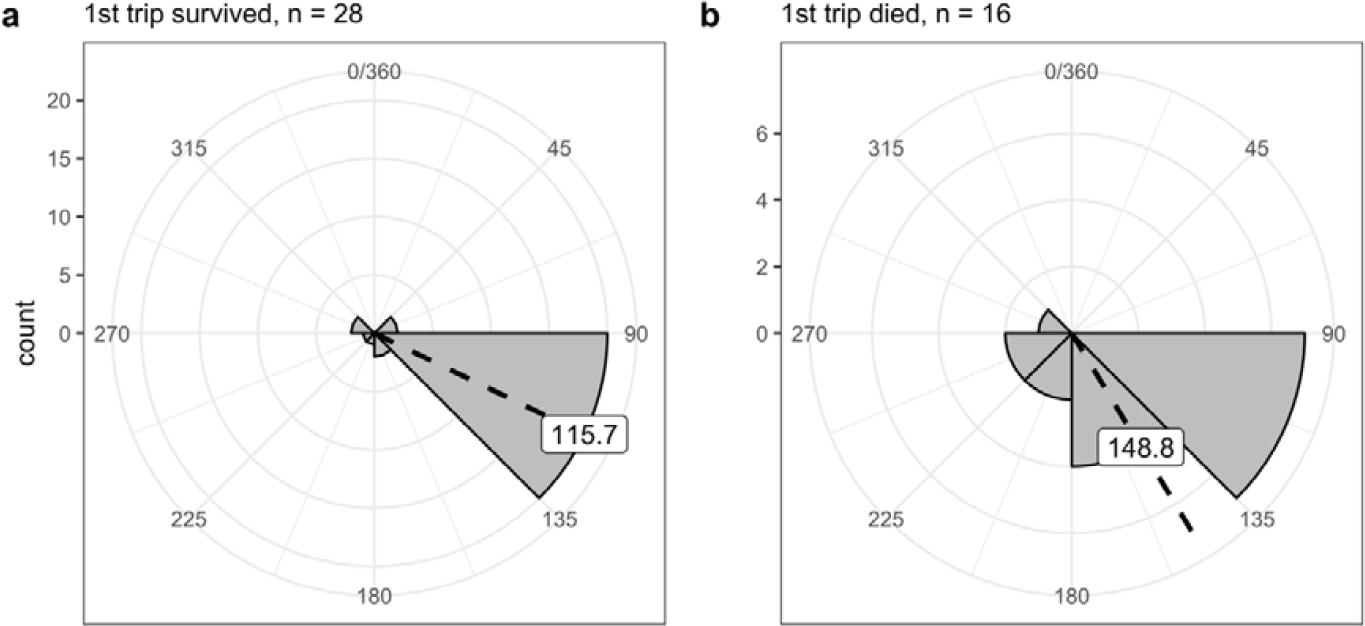
Circular histograms of dispersal direction for pups that survived vs died on their first foraging trip. The dashed line and label is the mean bearing.

**Figure 5.**
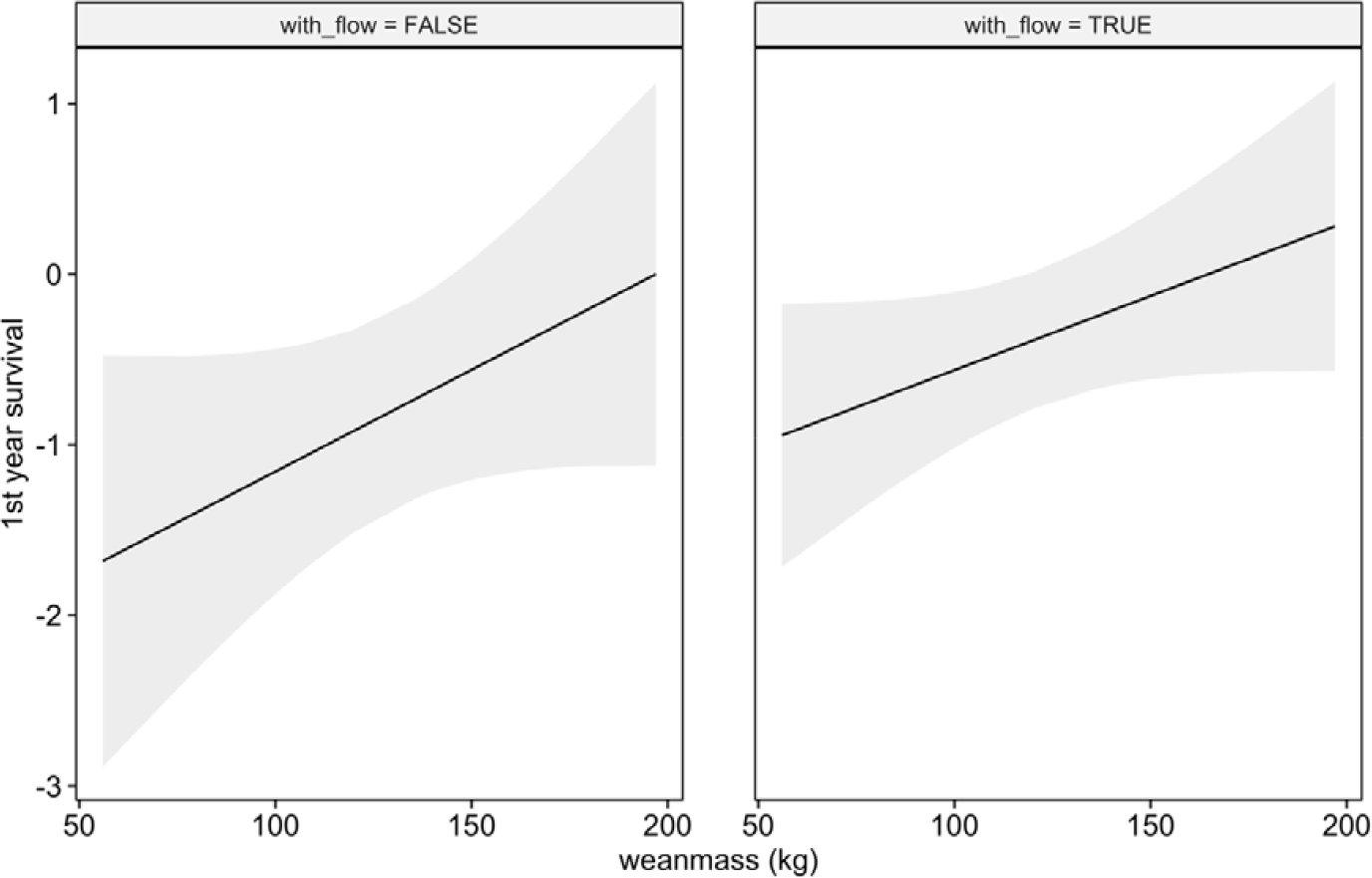
Predicted effects of how travelling in the east-southeast direction and wean mass affects first year survival of southern elephant seal weaners. Solid line = predicted fit; grey shaded band = SE.

## Discussion

Juvenile survival is a key determinant of population trajectory, particularly in long-lived slowly reproducing species like elephant seals [1]. We investigated the foraging movement of naïve, newly weaned elephant seal pups and related this to ocean currents and their subsequent survival. We found that travelling with the flow and being heavier at weaning are important factors that improve first-year survival rates. Additionally, dispersal direction from the colony was different between weaners and adult females, suggesting that dispersal behaviours can change over time and with experience. In other words, seals can manage their maiden trip with minimal information that is either inherited or learned. This is because simply going with the flow gets them to areas which improve their odds of survival.

### Dispersal direction and survival of weaners

Weaners travelling within the predominant east-southeast current demonstrated a 1.5 times greater survival rate compared to those moving westward against the current. This observation contrasts with findings on Kerguelen juveniles which showed no difference in dispersal direction between survivors and non-survivors [6]. The disparity might stem from the broader directional range (southeast, 90-degree range) used in that study’s analysis [6] relative to the range in this study (east-southeast, 45-degree range).

Previous studies have shown that southern elephant seal pups from Macquarie Island [21,51] and the Kerguelen Islands [6,23] generally disperse south-eastward from their birthplaces, aligning with the dominant ocean currents in those areas. In contrast, pups from King George Island (South Shetland Islands) predominantly travel south-westward from their colonies [26]. Meanwhile, pups from Marion Island exhibit no fixed dispersal directions [52]. These varied patterns could be attributed to the distinct bathymetric and hydrological characteristics surrounding each colony. For example, the region around King George Island is a lot closer to Antarctica and is less open compared to the other colonies [26]. In contrast, Marion Island pups may be more influenced by smaller scale currents and frontal structures than larger ocean currents like the eastward flowing ACC [52].

The seal tracks diverging from the particle trace during their foraging journey indicate that they are not merely drifting with the current. Instead, it appears while weaners swim without a set destination, they are influenced by the dominant ACC, steering them primarily south-eastward. As they travel, young seals may make subtle directional shifts [53] based on cues such as temperature gradients, currents, topography and olfactory signals produced by dimethyl sulphide from phytoplankton [54–57]. Nonetheless, the prevailing current ensures a consistent south-easterly trajectory. Seals that deviate from this general behaviour might be influenced by specific environmental or behavioural factors that counteract the current’s influence. However, it is interesting to question why these factors would result in poorer survival.

### Oceanic currents as a passive transport mechanism

In the first weeks of life, southern elephant seal pups are least experienced and rely heavily on the energy stores given to them by their mothers [58]. Swimming consumes a lot of energy for mammals [59,60]. Therefore, swimming with the flow of dominant ocean currents initially benefits the pups; it allows them to cover greater distances without using more energy [61,62]. It is particularly advantageous for bigger weaners, who can leverage their energy reserves, allowing them more time to find food before their stores deplete. However, it is important to consider that swimming downstream is a short-term advantage. To return to their colonies, the pups eventually must swim against these currents, which could negate some of the energy they initially saved.

Passive transportation downstream provides pups with two potential benefits. Firstly, it helps them move quickly away from areas near their colonies where predators like sharks and killer whales are likely to be found, thus reducing their risk of predation [6,63,64]. Secondly, it facilitates opportunistic exploration across diverse oceanic regions where they can discover and exploit prey patches (Field et al., 2005). The natural flow of currents can inadvertently steer seals towards zones where convergent surface currents result in the concentration of prey. For example, female southern elephant seals from Kerguelen typically swim downstream, to the east of the island, where they come across a greater number of eddies [65]. These oceanic features play a role in clustering prey into denser aggregations, vital for seals and other apex marine predators during their foraging excursions [65–68]. In contrast, juveniles that forage in areas with weak eddy activity face higher chances of mortality [6]. Observations of dispersal patterns among different southern elephant seal colonies suggest that the downstream-upstream (east-west) variation in prey distribution could be a widespread feature surrounding Southern Ocean islands.

### Differences in pups and adult females dispersal

Given their physiological and size constraints, younger seals do not travel as far as older seals on their foraging trips [25,69]. However, with the assistance of downstream currents, they can effectively navigate through the polar frontal zone to eddy-rich areas [70,71]. Notably, a yearling managed to cover approximately 5200 km eastwards from Macquarie Island to Peter the First Island, likely aided by downstream swimming [72].

Adult female seals display dispersal patterns that diverge significantly from those of weaned pups. This distinction indicates that ocean currents may initially lead young seals to adequate foraging areas, yet these areas may not be the most advantageous. The choice of habitat is the result of a complex interplay between life history strategies, environmental conditions, and predation risks, rather than just body size or energy maximisation [73]. In the critical early stages of life, seal pups develop essential diving and foraging skills [74] that are crucial for their long-term survival [75]. As they learn from experience, their random movements become more refined and directed, enabling them to reach more optimal foraging areas. This is shown by the behaviour of experienced adult females, which tend to travel more directly and further south towards Antarctic waters [25,42] that are more productive [76]. As seals mature, they are likely to discover superior foraging grounds to which they develop a strong fidelity [77–79].

### Body weight and dispersal effects on juvenile survival

Body weight at weaning is a key factor in the survival rates of juvenile seals. However, this relationship is not linear but best described by a quadratic relation, indicating a cost to being either too big or too small [80]. Generally, heavier weaners are more likely to survive their first year [18,81]. But weight alone does not guarantee survival. For example, some lighter seals that swam with the current also survived their first year. To illustrate the unpredictable nature of survival, consider two contrasting cases. One seal (mq1-17217-95), despite being the lightest, survived its first year, while another (mq4-cleo-00), being the heaviest, died during its first foraging trip, possibly due to predation. These varied outcomes, even under similar conditions, suggest that survival is a complex interplay between predictable (‘deterministic’) and unpredictable (‘stochastic’) factors.

### Conclusion

Our study underscores that beyond weaning mass, the act of swimming downstream significantly affects the survival of newly independent southern elephant seal pups, who lack prior knowledge of productive feeding grounds. Swimming with the prevailing current offers a passive mode of travel that not only reduces the threat of predators near their birth colony but also potentially carries them to regions abundant with prey. As these juvenile seals become more seasoned, they begin to intentionally navigate away from the current towards chosen foraging destinations. Investigating the impact of the primary eastward current in the sub-Antarctic on the circumpolar patterns of movement for southern elephant seals, and perhaps other marine predators, could broaden our comprehension of Southern Ocean ecology.

## Acknowledgements

The study was carried out at Macquarie Island under ethics approval from the Australian Antarctic Animal Ethics Committee (AAS 2265 & AAS 2794) and the Tasmanian Parks and Wildlife Service. The seal tracking data were sourced from the Integrated Marine Observing System (IMOS). IMOS is supported by the Australian Government through the National Collaborative Research Infrastructure Strategy. Logistics to Macquarie Island were provided by the Australian Antarctic Division. We thank the expeditioners at Macquarie Island between 1993 and 2014 for helping us to mark, weigh, measure and resight marked seals.

## References

1. McMahon CR, Bester MN, Burton HR, Hindell MA, Bradshaw CJAA. 2005 Population status, trends and a re-examination of the hypotheses explaining the recent declines of the southern elephant seal Mirounga leonina. Mammal Rev. 35, 82–100. (doi:10.1111/j.1365-2907.2005.00055.x)

2. McMahon CR, Hindell MA, Burton HR, Bester MN. 2005 Comparison of southern elephant seal poulations, and observations of a population on a demographic knife-edge. Mar. Ecol. Prog. Ser. 288, 273–283. (doi:10.3354/meps288273)

3. Daunt F, Afanasyev V, Adam A, Croxall JP, Wanless S. 2007 From cradle to early grave: Juvenile mortality in European shags Phalacrocorax aristotelis results from inadequate development of foraging proficiency. Biol. Lett. 3. (doi:10.1098/rsbl.2007.0157)

4. Enstipp MR, Bost C-A, Le Bohec C, Bost C, Le Maho Y, Weimerskirch H, Handrich Y. 2017 Apparent changes in body insulation of juvenile king penguins suggest an energetic challenge during their early life at sea. J. Exp. Biol. 220, 2666–2678. (doi:10.1242/jeb.160143)

5. Harel R, Horvitz N, Nathan R. 2016 Adult vultures outperform juveniles in challenging thermal soaring conditions. Sci. Rep. 6, 27865. (doi:10.1038/srep27865)

6. Cox SL, Authier M, Orgeret F, Weimerskirch H, Guinet C. 2020 High mortality rates in a juvenile free-ranging marine predator and links to dive and forage ability. Ecol. Evol. 10, 410–430. (doi:10.1002/ece3.5905)

7. Gomes A, Pereira J, Bugoni L. 2009 Age-specific diving and foraging behavior of the great grebe (podicephorus major). Waterbirds 32, 149–156. (doi:10.1675/063.032.0118)

8. Beauplet G, Barbraud C, Chambellant M, Guinet C. 2005 Interannual variation in the post-weaning and juvenile survival of subantarctic fur seals: Influence of pup sex, growth rate and oceanographic conditions. J. Anim. Ecol. 74. (doi:10.1111/j.1365-2656.2005.01016.x)

9. Estes JA, Riedman ML, Staedler MM, Tinker MT, Lyon BE. 2003 Individual variation in prey selection by sea otters: patterns, causes and implications. J. Anim. Ecol. 72, 144–155. (doi:10.1046/j.1365-2656.2003.00690.x)

10. Fowler SL, Costa DP, Arnould JPY. 2007 Ontogeny of movements and foraging ranges in the australian sea lion. Mar. Mammal Sci. 23, 598–614. (doi:10.1111/J.1748-7692.2007.00134.X)

11. Berger V, Reichert S, Lahdenperä M, Jackson J, Htut W, Lummaa V. 2021 The elephant in the family: Costs and benefits of elder siblings on younger offspring life-history trajectory in a matrilineal mammal. J. Anim. Ecol. 90, 2663–2677. (doi:10.1111/1365-2656.13573)

12. Lamon N, Neumann C, Gruber T, Zuberbühler K. 2017 Kin-based cultural transmission of tool use in wild chimpanzees. Sci. Adv. 3, e1602750. (doi:10.1126/sciadv.1602750)

13. Bender CE, Herzing DL, Bjorklund DF. 2009 Evidence of teaching in atlantic spotted dolphins (Stenella frontalis) by mother dolphins foraging in the presence of their calves. Anim. Cogn. 12, 43–53. (doi:10.1007/S10071-008-0169-9/TABLES/2)

14. Bowen WD, Boness DJ, Iverson SJ. 2011 Diving behaviour of lactating harbour seals and their pups during maternal foraging trips. Httpsdoiorg101139z99-065 77, 978–988. (doi:10.1139/Z99-065)

15. Gjertz I, Kovacs KM, Lydersen C Wiig. 2000 Movements and diving of bearded seal (Erignathus barbatus) mothers and pups during lactation and post-weaning. Polar Biol. 23, 559–566. (doi:10.1007/S003000000121/METRICS)

16. Sato K, Mitani Y, Kusagaya H, Naito Y. 2003 Synchronous shallow dives by weddell seal mother-pup pairs during lactation. Mar. Mammal Sci. 19, 384–395. (doi:10.1111/J.1748-7692.2003.TB01116.X)

17. Hindell MA, McConnell BJ, Fedak MA, Slip DJ, Burton HR, Reijnders PJHH, McMahon CR. 1999 Environmental and physiological determinants of successful foraging by naive southern elephant seal pups during their first trip to sea. Can. J. Zool. 77, 1807–1821. (doi:10.1139/z99-154)

18. McMahon CR, Burton HR, Bester MN. 2000 Weaning mass and the future survival of juvenile southern elephant seals, Mirounga leonina, at Macquarie Island. Antarct. Sci. 12, 149–153. (doi:10.1017/s0954102000000195)

19. McMahon CR, Harcourt RG, Burton HR, Daniel O, Hindell MA. 2017 Seal mothers expend more on offspring under favourable conditions and less when resources are limited. J. Anim. Ecol. 86, 359–370. (doi:10.1111/1365-2656.12611)

20. McMahon CR, New LF, Fairley EJ, Hindell MA, Burton HR. 2015 The effects of body size and climate on post-weaning survival of elephant seals at Heard Island. J. Zool. 297, 301–308. (doi:10.1111/jzo.12279)

21. McConnell B, Fedak M, Burton HR, Engelhard GH, Reijnders PJH. 2002 Movements and foraging areas of naïve, recently weaned southern elephant seal pups. J. Anim. Ecol. 71, 65–78. (doi:10.1046/j.0021-8790.2001.00576.x)

22. de Grissac S, Bartumeus F, Cox SL, Weimerskirch H. 2017 Early-life foraging: Behavioral responses of newly fledged albatrosses to environmental conditions. Ecol. Evol. 7, 6766–6778. (doi:10.1002/ece3.3210)

23. Orgeret F, Cox SL, Weimerskirch H, Guinet C. 2019 Body condition influences ontogeny of foraging behavior in juvenile southern elephant seals. Ecol. Evol. 9, 223–236. (doi:10.1002/ECE3.4717)

24. Campagna J, Lewis MN, González Carman V, Campagna C, Guinet C, Johnson M, Davis RW, Rodríguez DH, Hindell MA. 2021 Ontogenetic niche partitioning in southern elephant seals from Argentine Patagonia. Mar. Mammal Sci. 37, 631–651. (doi:10.1111/mms.12770)

25. Field IC, Bradshaw CJA, Burton HR, Sumner MD, Hindell MA. 2005 Resource partitioning through oceanic segregation of foraging juvenile southern elephant seals (Mirounga leonina). Oecologia 142, 127–135. (doi:10.1007/s00442-004-1704-2)

26. Bornemann H, Kreyscher M, Ramdohr S, Martin T, Carlini A, Sellmann L, Plötz J. 2000 Southern elephant seal movements and Antarctic sea ice. Antarct. Sci. 12, 3–15. (doi:10.1017/s095410200000002x)

27. Riotte-Lambert L, Weimerskirch H. 2013 Do naive juvenile seabirds forage differently from adults? Proc. R. Soc. B Biol. Sci. 280, 20131434. (doi:10.1098/rspb.2013.1434)

28. Ling JK, Bryden MM. 1992 Mirounga leonina. Mamm. Species, 1–8. (doi:10.2307/3504169)

29. Hindell MA, Bradshaw CJA, Sumner MD, Michael KJ, Burton HR. 2003 Dispersal of female southern elephant seals and their prey consumption during the austral summer: relevance to management and oceanographic zones. J. Appl. Ecol. 40, 703–715. (doi:10.1046/j.1365-2664.2003.00832.x)

30. Lübcker N, Reisinger RR, Oosthuizen WC, Bruyn PJN de, Tonder A van, Pistorius PA, Bester MN. 2017 Low trophic level diet of juvenile southern elephant seals Mirounga leonina from Marion Island: a stable isotope investigation using vibrissal regrowths. Mar. Ecol. Prog. Ser. 577, 237–250. (doi:10.3354/meps12240)

31. Newland C, Field IC, Cherel Y, Guinet C, Bradshaw CJA, Harcourt R, Hindell MA. 2011 Diet of juvenile southern elephant seals reappraised by stable isotopes in whiskers. Mar. Ecol. Prog. Ser. 424. (doi:10.3354/meps08769)

32. Walters A, Lea MA, van den Hoff J, Field IC, Virtue P, Sokolov S, Pinkerton MH, Hindell MA. 2014 Spatially explicit estimates of prey consumption reveal a new krill predator in the Southern Ocean. PLoS ONE 9, e86452. (doi:10.1371/journal.pone.0086452)

33. Hindell MA et al. 2016 Circumpolar habitat use in the southern elephant seal: Implications for foraging success and population trajectories. Ecosphere 7, e01213. (doi:10.1002/ecs2.1213)

34. Chapman CC, Lea M-A, Meyer A, Sallée J-B, Hindell M. 2020 Defining Southern Ocean fronts and their influence on biological and physical processes in a changing climate. Nat. Clim. Change 10, 209–219. (doi:10.1038/s41558-020-0705-4)

35. Gordine SA, Fedak MA, Boehme L. 2019 The importance of Southern Ocean frontal systems for the improvement of body condition in southern elephant seals. Aquat. Conserv. Mar. Freshw. Ecosyst. 29, 283–304. (doi:10.1002/aqc.3183)

36. Pyke GH, Pulliam HR, Charnov EL. 1977 Optimal Foraging - A Selective Review of Theory and Tests. Q. Rev. Biol. 52, 137–154. (doi:10.1086/409852)

37. Clarke J, Kerry K, Fowler CE, Lawless R, Eberhard SM, Murphy R. 2003 Post-fledging and winter migration of Adélie penguins Pygoscelis adeliae in the Mawson region of East Antarctica. Mar. Ecol. Prog. Ser. 248. (doi:10.3354/meps248267)

38. Ream RR, Sterling JT, Loughlin TR. 2005 Oceanographic features related to northern fur seal migratory movements. Deep Sea Res. Part II Top. Stud. Oceanogr. 52, 823–843. (doi:10.1016/j.dsr2.2004.12.021)

39. Thiebot J-B, Lescroël A, Barbraud C, Bost C-A. 2013 Three-dimensional use of marine habitats by juvenile emperor penguins Aptenodytes forsteri during post-natal dispersal. Antarct. Sci. 25, 536–544. (doi:10.1017/S0954102012001198)

40. Hindell MA, Slip DJ, Burton HR. 1991 The diving behaviour of adult male and female southern elephant seals, Mirounga leonina (Pinnipedia: Phocidae). Aust. J. Zool. 39, 499–508. (doi:10.1071/ZO9910595)

41. McMahon CR, Burton HR, Bester MN. 1999 First-year survival of southern elephant seals, Mirounga leonina, at sub-Antarctic Macquarie Island. Polar Biol. 21, 279–284. (doi:10.1007/S003000050363/METRICS)

42. Hindell MA, Sumner M, Bestley S, Wotherspoon S, Harcourt RG, Lea MA, Alderman R, McMahon CR. 2017 Decadal changes in habitat characteristics influence population trajectories of southern elephant seals. Glob. Change Biol. 23, 5136–5150. (doi:10.1111/gcb.13776)

43. Foo D, McMahon C, Hindell M, Goldsworthy S, Bailleul F. 2019 Influence of shelf oceanographic variability on alternate foraging strategies in long-nosed fur seals. Mar. Ecol. Prog. Ser. 615, 189–204. (doi:10.3354/meps12922)

44. Jonsen ID et al. 2020 A continuous-time state-space model for rapid quality control of argos locations from animal-borne tags. Mov. Ecol. 8. (doi:10.1186/s40462-020-00217-7)

45. Jonsen ID, Grecian WJ, Phillips L, Carroll G, McMahon C, Harcourt RG, Hindell MA, Patterson TA. 2023 aniMotum, an R package for animal movement data: Rapid quality control, behavioural estimation and simulation. Methods Ecol. Evol. 14, 806–816. (doi:10.1111/2041-210X.14060)

46. Sumner M. 2021 Currently.

47. Agostinelli C, Lund U. 2023 Circular Statistics.

48. Landler L, Ruxton GD, Malkemper EP. 2021 Advice on comparing two independent samples of circular data in biology. Sci. Rep. 11, 1–10. (doi:10.1038/s41598-021-99299-5)

49. Bartoń K. 2009 MuMIn: Multi-model inference.

50. Hartig F. 2022 DHARMa - Residual Diagnostics for HierARchical Models.

51. van den Hoff J. 2002 Migrations of Juvenile Southern Elephant Seals from Macquarie Island. University of Tasmania.

52. McIntyre T, Oosthuizen WC, Bester MN, Hindell MA, Reisinger RR, Tosh CA, van den Hoff J, de Bruyn PJN. 2023 Tracking the foraging migrations of Marion Island southern elephant seals (Mirounga leonina) during their first year of life. Mar. Mammal Sci., 1–19. (doi:10.1111/mms.13078)

53. Painter KJ, Hillen T. 2015 Navigating the flow: individual and continuum models for homing in flowing environments. J. R. Soc. Interface 12, 20150647. (doi:10.1098/rsif.2015.0647)

54. Luschi P. 2013 Long-distance animal migrations in the oceanic environment: orientation and navigation correlates. Int. Sch. Res. Not. 2013, e631839. (doi:10.1155/2013/631839)

55. Matsumura M, Watanabe YY, Robinson PW, Miller PJO, Costa DP, Miyazaki N. 2011 Underwater and surface behavior of homing juvenile northern elephant seals. J. Exp. Biol. 214, 629–636. (doi:10.1242/jeb.048827)

56. Mattern T, Ellenberg U, Houston DM, Lamare M, Davis LS, Heezik Y van, Seddon PJ. 2013 Straight line foraging in yellow-eyed penguins: new insights into cascading fisheries effects and orientation capabilities of marine predators. PLOS ONE 8, e84381. (doi:10.1371/journal.pone.0084381)

57. Tosh C, Steyn J, Bornemann H, Van Den Hoff J, Stewart B, Plötz J, Bester M. 2012 Marine habitats of juvenile southern elephant seals from Marion Island. Aquat. Biol. 17, 71–79. (doi:10.3354/ab00463)

58. Bryden MM. 1968 Growth and function of the subcutaneous fat of the elephant seal. Nature 220, 597–599. (doi:10.1038/220597a0)

59. Miller PJO, Biuw M, Watanabe YY, Thompson D, Fedak MA. 2012 Sink fast and swim harder! Round-trip cost-of-transport for buoyant divers. J. Exp. Biol. 215, 3622–3630. (doi:10.1242/jeb.070128)

60. Williams TM, Davis RW, Fuiman LA, Francis J, Le Boeuf BL, Horning M, Calambokidis J, Croll DA. 2000 Sink or swim: strategies for cost-efficient diving by marine mammals. Science 288, 133–136.

61. Lea MA, Johnson D, Ream R, Sterling J, Melin S, Gelatt T. 2009 Extreme weather events influence dispersal of naive northern fur seals. Biol. Lett. 5, 252–257. (doi:10.1098/rsbl.2008.0643)

62. Weimerskirch H, Guionnet T, Martin J, Shaffer SA, Costa DP. 2000 Fast and fuel efficient? Optimal use of wind by flying albatrosses. Proc. R. Soc. Lond. B Biol. Sci. 267, 1869–1874. (doi:10.1098/rspb.2000.1223)

63. Henderson AF, McMahon CR, Harcourt R, Guinet C, Picard B, Wotherspoon S, Hindell MA. 2020 Inferring variation in southern elephant seal at-sea mortality by modelling tag failure. Front. Mar. Sci. 7. (doi:10.3389/fmars.2020.517901)

64. Horning M, Mellish JAE. 2012 Predation on an upper trophic marine predator, the steller sea lion: Evaluating high juvenile mortality in a density dependent conceptual framework. PLoS ONE 7. (doi:10.1371/journal.pone.0030173)

65. Green DB, Bestley S, Trebilco R, Corney S, Lehodey P, McMahon C, Guinet C, Hindell MA. 2020 Modelled mid-trophic pelagic prey fields improve understanding of marine predator foraging behaviour. Ecography. (doi:10.1111/ecog.04939)

66. Cotte C et al. 2015 Flexible preference of southern elephant seals for distinct mesoscale features within the Antarctic Circumpolar Current. Prog. Oceanogr. 131, 46–58. (doi:10.1016/j.pocean.2014.11.011)

67. Fahlbusch JA, Czapanskiy MF, Calambokidis J, Cade DE, Abrahms B, Hazen EL, Goldbogen JA. 2022 Blue whales increase feeding rates at fine-scale ocean features. Proc. R. Soc. B 289. (doi:10.1098/RSPB.2022.1180)

68. Oliver MJ et al. 2019 Central place foragers select ocean surface convergent features despite differing foraging strategies. Sci. Rep. 9, 157. (doi:10.1038/s41598-018-35901-7)

69. Irvine LG, Hindell MA, van den Hoff J, Burton HR. 2000 The influence of body size on dive duration of underyearling southern elephant seals (Mirounga leonina). J. Zool. 251, 463–471. (doi:10.1017/S0952836900008062)

70. Bailleul F, Cotté C, Guinet C. 2010 Mesoscale eddies as foraging area of a deep-diving predator, the southern elephant seal. Mar. Ecol. Prog. Ser. 408, 251–264. (doi:10.3354/meps08560)

71. McConnell BJ, Chambers C, Fedak MA. 1992 Foraging ecology of southern elephant seals in relation to the bathymetry and productivity of the southern-ocean. Antarct. Sci. 4, 393–398.

72. Hindell MA, Mcmahon CR. 2000 Long distance movement of a southern elephant seal (mirounga leonina) from macquarie island to peter 1 oy. Mar. Mammal Sci. 16, 504–507. (doi:10.1111/j.1748-7692.2000.tb00944.x)

73. Hindell MA, McMahon CR, Jonsen I, Harcourt R, Arce F, Guinet C. 2021 Inter- and intrasex habitat partitioning in the highly dimorphic southern elephant seal. Ecol. Evol. 11, 1620–1633. (doi:10.1002/ece3.7147)

74. Zeppelin T, Pelland N, Sterling J, Brost B, Melin S, Johnson D, Lea M-A, Ream R. 2019 Migratory strategies of juvenile northern fur seals (Callorhinus ursinus): bridging the gap between pups and adults. Sci. Rep. 9, 13921. (doi:10.1038/s41598-019-50230-z)

75. Baylis AMM, Page B, Peters K, McIntosh R, McKenzie J, Goldsworthy S. 2005 The ontogeny of diving behaviour in New Zealand fur seal pups (Arctocephalus forsteri). Can. J. Zool. 83, 1149–1161. (doi:10.1139/z05-097)

76. Thomalla SJ, Nicholson S-A, Ryan-Keogh TJ, Smith ME. 2023 Widespread changes in Southern Ocean phytoplankton blooms linked to climate drivers. Nat. Clim. Change 13, 975–984. (doi:10.1038/s41558-023-01768-4)

77. Authier M, Bentaleb I, Ponchon A, Martin C, Guinet C. 2012 Foraging fidelity as a recipe for a long life: Foraging strategy and longevity in male southern elephant seals. PLoS ONE 7, e32026. (doi:10.1371/journal.pone.0032026)

78. Bradshaw CJA, Hindell MA, Sumner MD, Michael KJ. 2004 Loyalty pays: Potential life history consequences of fidelity to marine foraging regions by southern elephant seals. Anim. Behav. 68, 1349–1360. (doi:10.1016/j.anbehav.2003.12.013)

79. Robinson PA et al. 2012 Foraging Behavior and Success of a Mesopelagic Predator in the Northeast Pacific Ocean: Insights From a Data-Rich Species, the Northern Elephant Seal. Plos One. (doi:10.1371/journal.pone.0036728)

80. McMahon CR, Burton HR, Bester MN. 2003 A demographic comparison of two southern elephant seal populations. J. Anim. Ecol. 72, 61–74. (doi:10.1046/j.1365-2656.2003.00685.x)

81. Oosthuizen WC, Altwegg R, Nevoux M, Bester MN, de Bruyn PJN. 2018 Phenotypic selection and covariation in the life-history traits of elephant seals: heavier offspring gain a double selective advantage. Oikos 127, 875–889. (doi:10.1111/oik.04998)

