## Supplementary Information 1 for "Swimming with the current improves juvenile survival in southern elephant seals"

### Supplementary material

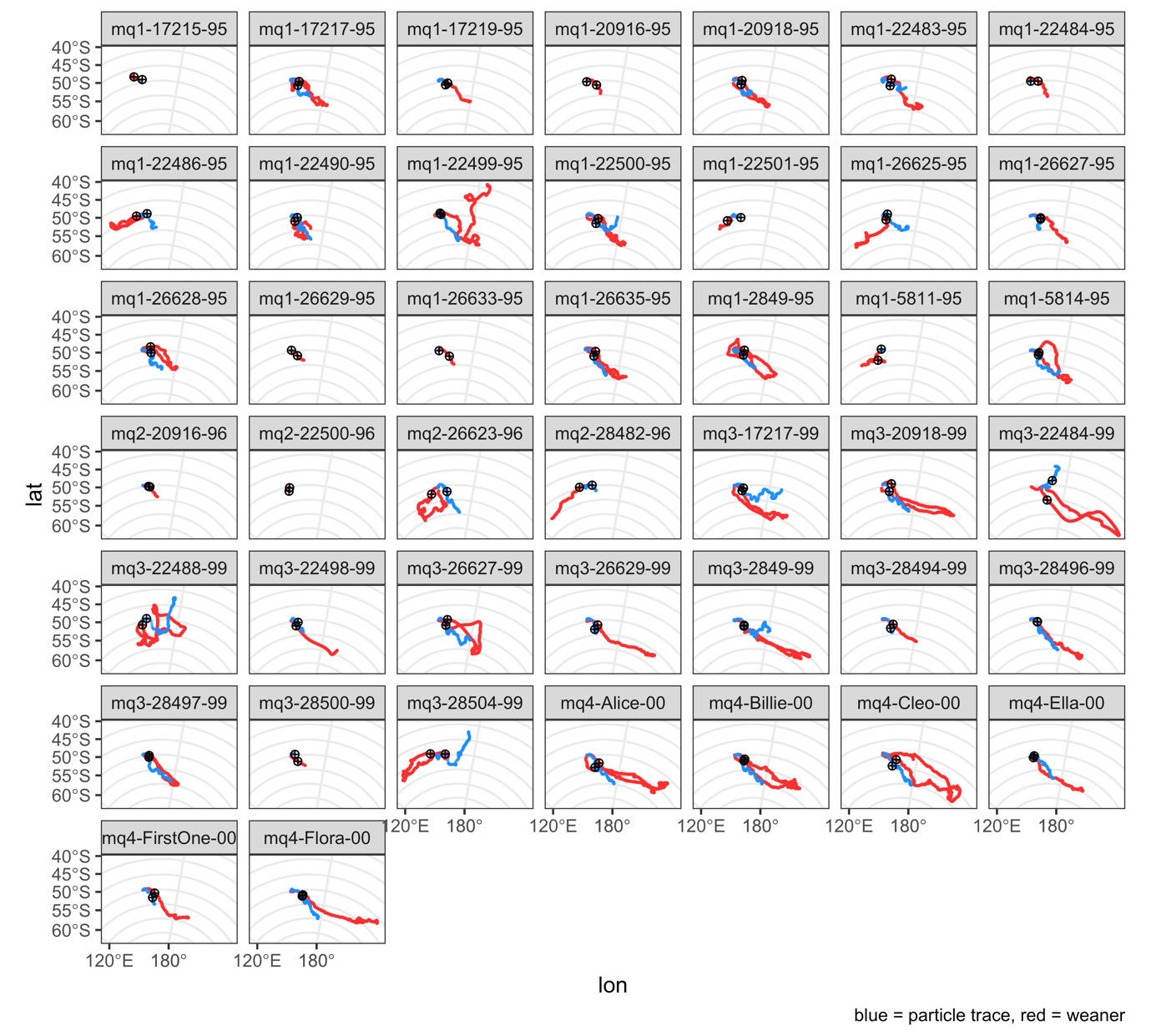

Supplementary Figure 1. First foraging trip tracks of weaners (red) and their corresponding particle traces (blue). The circular shapes represent the location what was used to calculate dispersal bearings.

Supplementary Table 1. Calculated dispersal bearing from Macquarie Island based on a reference location for weaners, adult females, and the particle traces. Reference location was the location on day 5 of the trip for weaners and adult females, whereas for particle traces it was based on the location that was closest to the corresponding weaners reference locations’ distance from the colony. Geolocation light loggers, gls.

| **Id** | **group** | **type** | **bearing (deg)** | **refence location distance to colony (km)** | **Start date** | **End date** | **trip duration (d)** | **compass direction** |
| --- | --- | --- | --- | --- | --- | --- | --- | --- |
| mq1-17215-95 | weaner | argos | 308.4 | 335.3 | 4/12/1995 | 26/12/1995 | 22 | W-NW |
| mq1-17217-95 | weaner | argos | 104.0 | 344.9 | 1/12/1995 | 9/4/1996 | 130 | E-SE |
| mq1-17219-95 | weaner | argos | 112.7 | 379.3 | 3/12/1995 | 8/1/1996 | 36 | E-SE |
| mq1-20916-95 | weaner | argos | 121.2 | 428.1 | 1/12/1995 | 29/12/1995 | 28 | E-SE |
| mq1-20918-95 | weaner | argos | 97.0 | 310.5 | 29/11/1995 | 28/4/1996 | 151 | E-SE |
| mq1-22483-95 | weaner | argos | 90.1 | 364.7 | 29/11/1995 | 4/4/1996 | 127 | E-SE |
| mq1-22484-95 | weaner | argos | 102.0 | 314.8 | 29/11/1995 | 21/12/1995 | 22 | E-SE |
| mq1-22486-95 | weaner | argos | 277.5 | 206.7 | 28/11/1995 | 26/3/1996 | 119 | W-NW |
| mq1-22490-95 | weaner | argos | 149.0 | 279.2 | 28/11/1995 | 28/4/1996 | 152 | SE-S |
| mq1-22499-95 | weaner | argos | 50.7 | 128.4 | 5/12/1995 | 2/6/1996 | 180 | NE-E |
| mq1-22500-95 | weaner | argos | 112.9 | 474.5 | 27/11/1995 | 3/5/1996 | 158 | E-SE |
| mq1-22501-95 | weaner | argos | 243.2 | 300.7 | 4/12/1995 | 18/1/1996 | 45 | SW-W |
| mq1-26625-95 | weaner | argos | 143.7 | 221.2 | 28/11/1995 | 22/3/1996 | 115 | SE-S |
| mq1-26627-95 | weaner | argos | 110.8 | 422.5 | 29/11/1995 | 11/2/1996 | 74 | E-SE |
| mq1-26628-95 | weaner | argos | 76.7 | 347.2 | 29/11/1995 | 5/4/1996 | 128 | NE-E |
| mq1-26629-95 | weaner | argos | 134.9 | 340.7 | 1/12/1995 | 23/12/1995 | 22 | E-SE |
| mq1-26633-95 | weaner | argos | 126.6 | 481.3 | 28/11/1995 | 19/12/1995 | 21 | E-SE |
| mq1-26635-95 | weaner | argos | 105.3 | 364.5 | 28/11/1995 | 13/5/1996 | 167 | E-SE |
| mq1-2849-95 | weaner | argos | 98.4 | 398.3 | 1/12/1995 | 15/4/1996 | 136 | E-SE |
| mq1-5811-95 | weaner | argos | 212.1 | 389.7 | 4/12/1995 | 27/1/1996 | 54 | S-SW |
| mq1-5814-95 | weaner | argos | 126.0 | 364.0 | 3/12/1995 | 4/5/1996 | 153 | E-SE |
| mq2-20916-96 | weaner | argos | 110.9 | 301.1 | 1/12/1996 | 12/12/1996 | 11 | E-SE |
| mq2-22500-96 | weaner | argos | 199.6 | 219.7 | 3/12/1996 | 24/12/1996 | 21 | S-SW |
| mq2-26623-96 | weaner | argos | 226.9 | 407.8 | 3/12/1996 | 10/6/1997 | 189 | SW-W |
| mq2-28482-96 | weaner | argos | 264.6 | 243.8 | 4/12/1996 | 2/2/1997 | 60 | SW-W |
| mq3-17217-99 | weaner | argos | 133.3 | 369.7 | 27/12/1999 | 27/6/2000 | 183 | E-SE |
| mq3-20918-99 | weaner | argos | 91.0 | 367.9 | 1/12/1999 | 24/5/2000 | 175 | E-SE |
| mq3-22484-99 | weaner | argos | 140.1 | 866.4 | 17/12/1999 | 21/5/2000 | 156 | SE-S |
| mq3-22488-99 | weaner | argos | 185.5 | 177.3 | 23/11/1999 | 16/8/2000 | 267 | S-SW |
| mq3-22498-99 | weaner | argos | 143.5 | 314.3 | 19/11/1999 | 17/1/2000 | 59 | SE-S |
| mq3-26627-99 | weaner | argos | 97.1 | 357.8 | 3/12/1999 | 12/6/2000 | 192 | E-SE |
| mq3-26629-99 | weaner | argos | 122.3 | 480.4 | 3/12/1999 | 9/4/2000 | 128 | E-SE |
| mq3-2849-99 | weaner | argos | 126.2 | 435.3 | 27/12/1999 | 27/6/2000 | 183 | E-SE |
| mq3-28494-99 | weaner | argos | 120.2 | 456.5 | 13/12/1999 | 1/2/2000 | 50 | E-SE |
| mq3-28496-99 | weaner | argos | 110.3 | 300.1 | 22/12/1999 | 28/11/2000 | 342 | E-SE |
| mq3-28497-99 | weaner | argos | 104.6 | 268.9 | 3/12/1999 | 13/9/2000 | 285 | E-SE |
| mq3-28500-99 | weaner | argos | 139.6 | 381.6 | 4/12/1999 | 26/12/1999 | 22 | SE-S |
| mq3-28504-99 | weaner | argos | 290.4 | 282.7 | 27/12/1999 | 19/1/2001 | 389 | W-NW |
| mq4-Alice-00 | weaner | argos | 131.4 | 583.7 | 24/12/2000 | 5/1/2002 | 377 | E-SE |
| mq4-Billie-00 | weaner | argos | 121.1 | 441.4 | 9/12/2000 | 29/8/2001 | 263 | E-SE |
| mq4-Cleo-00 | weaner | argos | 118.1 | 571.4 | 8/12/2000 | 18/8/2001 | 253 | E-SE |
| mq4-Ella-00 | weaner | argos | 135.3 | 176.9 | 24/12/2000 | 13/4/2001 | 110 | SE-S |
| mq4-FirstOne-00 | weaner | argos | 114.7 | 489.8 | 23/11/2000 | 30/1/2001 | 68 | E-SE |
| mq4-Flora-00 | weaner | argos | 121.3 | 521.0 | 24/12/2000 | 26/6/2001 | 184 | E-SE |
| b131pm_01 | adult female | gls | 157.5 | 393.8 | 10/2/2001 | 29/9/2001 | 231 | SE-S |
| b143pm_04 | adult female | gls | 154.9 | 557.7 | 17/2/2004 | 7/10/2004 | 233 | SE-S |
| b279pm_02 | adult female | gls | 171.6 | 537.5 | 2/2/2002 | 21/9/2002 | 231 | SE-S |
| b347pm_04 | adult female | gls | 231.2 | 369.1 | 2/2/2004 | 5/10/2004 | 246 | SW-W |
| b362pm_01 | adult female | gls | 161.6 | 654.1 | 21/2/2001 | 7/10/2001 | 228 | SE-S |
| b533pm_00 | adult female | gls | 221.2 | 441.3 | 3/2/2000 | 27/9/2000 | 237 | S-SW |
| b546pm_04 | adult female | gls | 151.7 | 391.3 | 14/2/2004 | 31/10/2004 | 260 | SE-S |
| b569pm_00 | adult female | gls | 180.2 | 581.4 | 3/3/2000 | 6/10/2000 | 217 | S-SW |
| b650pm_02 | adult female | gls | 238.2 | 308.6 | 6/2/2002 | 21/9/2002 | 227 | SW-W |
| b864pm_00 | adult female | gls | 148.1 | 432.0 | 14/2/2000 | 17/7/2000 | 154 | SE-S |
| b883pm_02 | adult female | gls | 148.5 | 360.3 | 1/2/2002 | 25/9/2002 | 236 | SE-S |
| b889pm_00 | adult female | gls | 154.9 | 388.9 | 6/2/2000 | 1/10/2000 | 238 | SE-S |
| b900pm_00 | adult female | gls | 180.2 | 543.9 | 25/2/2000 | 9/10/2000 | 227 | S-SW |
| b900pm_01 | adult female | gls | 147.1 | 466.5 | 20/2/2001 | 27/8/2001 | 188 | SE-S |
| b900pm_04 | adult female | gls | 161.7 | 251.8 | 14/2/2004 | 4/10/2004 | 233 | SE-S |
| c041pm_02 | adult female | gls | 92.0 | 135.1 | 3/2/2002 | 24/9/2002 | 233 | E-SE |
| c064pm_00 | adult female | gls | 150.3 | 293.6 | 21/2/2000 | 10/10/2000 | 232 | SE-S |
| c064pm_01 | adult female | gls | 165.0 | 306.9 | 17/2/2001 | 3/10/2001 | 228 | SE-S |
| c064pm_04 | adult female | gls | 161.3 | 340.1 | 25/2/2004 | 16/10/2004 | 234 | SE-S |
| c161pm_04 | adult female | gls | 221.9 | 185.2 | 15/2/2004 | 4/10/2004 | 232 | S-SW |
| c162pm_02 | adult female | gls | 129.0 | 488.3 | 19/2/2002 | 9/10/2002 | 232 | E-SE |
| c162pm_04 | adult female | gls | 142.4 | 554.0 | 15/2/2004 | 6/10/2004 | 234 | SE-S |
| c163pm_01 | adult female | gls | 30.7 | 503.2 | 5/2/2001 | 10/10/2001 | 247 | N-NE |
| c163pm_02 | adult female | gls | 30.7 | 503.2 | 5/2/2001 | 10/10/2001 | 247 | N-NE |
| c163pm_05 | adult female | gls | 30.7 | 503.2 | 5/2/2001 | 10/10/2001 | 247 | N-NE |
| c200pm_02 | adult female | gls | 149.5 | 522.7 | 6/2/2002 | 22/9/2002 | 228 | SE-S |
| c200pm_04 | adult female | gls | 136.8 | 438.1 | 11/2/2004 | 13/10/2004 | 245 | SE-S |
| c310pm_02 | adult female | gls | 89.6 | 104.1 | 5/2/2002 | 5/6/2002 | 120 | NE-E |
| c312pm_02 | adult female | gls | 318.0 | 95.2 | 3/2/2002 | 29/9/2002 | 238 | NW-N |
| c699pm_00 | adult female | gls | 186.7 | 583.4 | 5/2/2000 | 24/9/2000 | 232 | S-SW |
| c923pm_02 | adult female | gls | 147.6 | 447.8 | 1/2/2002 | 29/9/2002 | 240 | SE-S |
| ct2_9916_04 | adult female | argos | 144.8 | 449.0 | 18/2/2004 | 20/7/2004 | 153 | SE-S |
| ct2_9919_04 | adult female | argos | 161.7 | 504.8 | 5/2/2004 | 11/7/2004 | 157 | SE-S |
| ct2_9925_04 | adult female | argos | 196.2 | 342.0 | 1/2/2004 | 30/7/2004 | 180 | S-SW |
| ct6_10006_05 | adult female | argos | 214.7 | 428.8 | 11/2/2005 | 4/5/2005 | 82 | S-SW |
| ct6_10010_05 | adult female | argos | 198.9 | 481.7 | 8/2/2005 | 20/4/2005 | 71 | S-SW |
| ct6_10015_05 | adult female | argos | 211.0 | 373.6 | 4/2/2005 | 22/9/2005 | 230 | S-SW |
| ct6_10016_05 | adult female | argos | 153.4 | 475.5 | 18/2/2005 | 16/3/2005 | 26 | SE-S |
| ct6_10017_05 | adult female | argos | 206.4 | 596.6 | 1/2/2005 | 31/8/2005 | 211 | S-SW |
| ct6_10018_05 | adult female | argos | 172.9 | 427.3 | 14/2/2005 | 19/9/2005 | 217 | SE-S |
| ct6_10020_05 | adult female | argos | 223.8 | 374.5 | 12/2/2005 | 6/10/2005 | 236 | S-SW |
| ct6_10021_05 | adult female | argos | 218.4 | 561.0 | 11/2/2005 | 17/9/2005 | 218 | S-SW |
| ct6_9920_05 | adult female | argos | 149.6 | 516.9 | 20/2/2005 | 16/4/2005 | 55 | SE-S |
| ct6_9925_05 | adult female | argos | 146.4 | 508.5 | 23/2/2005 | 28/8/2005 | 186 | SE-S |
| f993pm_05 | adult female | gls | 132.6 | 192.1 | 14/2/2005 | 30/9/2005 | 228 | E-SE |
| h285pm_04 | adult female | gls | 221.6 | 278.9 | 18/2/2004 | 9/10/2004 | 234 | S-SW |
| h833pm_04 | adult female | gls | 138.6 | 235.9 | 17/2/2004 | 8/10/2004 | 234 | SE-S |
| ct64-M040-09 | adult female | argos | 196.7 | 236.1 | 2/2/2010 | 5/8/2010 | 184 | S-SW |
| ct64-M043-09 | adult female | argos | 207.1 | 635.0 | 1/2/2010 | 24/10/2010 | 265 | S-SW |
| ct64-M044-09 | adult female | argos | 172.4 | 421.6 | 8/2/2010 | 6/10/2010 | 240 | SE-S |
| ct64-M052-09 | adult female | argos | 196.4 | 389.7 | 10/2/2010 | 4/8/2010 | 175 | S-SW |
| ct64-M059-09 | adult female | argos | 204.2 | 493.9 | 4/2/2010 | 23/9/2010 | 231 | S-SW |
| ct64-M061-09 | adult female | argos | 146.6 | 418.3 | 15/2/2010 | 26/4/2010 | 70 | SE-S |
| ct64-M721-09 | adult female | argos | 147.0 | 497.5 | 4/2/2010 | 21/9/2010 | 229 | SE-S |
| ct64-M746-09 | adult female | argos | 128.1 | 604.1 | 4/2/2010 | 26/9/2010 | 234 | E-SE |
| ct64-M752-09 | adult female | argos | 141.3 | 423.2 | 4/2/2010 | 23/9/2010 | 231 | SE-S |
| ct64-M979-09 | adult female | argos | 192.3 | 591.1 | 1/2/2010 | 1/10/2010 | 242 | S-SW |
| ct64-M994-09 | adult female | argos | 146.8 | 345.5 | 5/2/2010 | 13/10/2010 | 250 | SE-S |
| mq1-17215-95 | particle trace |  | 18.9 | 52.7 | 15/12/1995 | 26/12/1995 | 11 | N-NE |
| mq1-17217-95 | particle trace |  | 130.0 | 340.6 | 6/12/1995 | 4/4/1996 | 120 | E-SE |
| mq1-17219-95 | particle trace |  | 127.7 | 323.2 | 7/12/1995 | 8/1/1996 | 32 | E-SE |
| mq1-20916-95 | particle trace |  | 140.3 | 44.8 | 19/12/1995 | 29/12/1995 | 10 | SE-S |
| mq1-20918-95 | particle trace |  | 125.9 | 304.2 | 6/12/1995 | 18/4/1996 | 134 | E-SE |
| mq1-22483-95 | particle trace |  | 130.4 | 366.9 | 6/12/1995 | 4/4/1996 | 120 | E-SE |
| mq1-22484-95 | particle trace |  | 113.0 | 42.6 | 29/11/1995 | 21/12/1995 | 22 | E-SE |
| mq1-22486-95 | particle trace |  | 77.9 | 204.0 | 1/12/1995 | 25/3/1996 | 115 | NE-E |
| mq1-22490-95 | particle trace |  | 114.0 | 284.6 | 13/12/1995 | 28/4/1996 | 137 | E-SE |
| mq1-22499-95 | particle trace |  | 77.8 | 127.9 | 10/12/1995 | 2/6/1996 | 175 | NE-E |
| mq1-22500-95 | particle trace |  | 137.7 | 479.5 | 3/12/1995 | 26/4/1996 | 145 | SE-S |
| mq1-22501-95 | particle trace |  | 114.8 | 275.7 | 13/12/1995 | 18/1/1996 | 36 | E-SE |
| mq1-26625-95 | particle trace |  | 84.8 | 223.3 | 5/12/1995 | 22/3/1996 | 108 | NE-E |
| mq1-26627-95 | particle trace |  | 117.3 | 430.8 | 9/12/1995 | 10/2/1996 | 63 | E-SE |
| mq1-26628-95 | particle trace |  | 113.7 | 349.1 | 30/11/1995 | 27/3/1996 | 118 | E-SE |
| mq1-26629-95 | particle trace |  | 90.0 | 58.0 | 13/12/1995 | 23/12/1995 | 10 | NE-E |
| mq1-26633-95 | particle trace |  | 105.2 | 45.0 | 10/12/1995 | 19/12/1995 | 9 | E-SE |
| mq1-26635-95 | particle trace |  | 134.8 | 369.4 | 6/12/1995 | 27/4/1996 | 143 | E-SE |
| mq1-2849-95 | particle trace |  | 125.7 | 396.7 | 2/12/1995 | 9/4/1996 | 129 | E-SE |
| mq1-5811-95 | particle trace |  | 356.3 | 55.3 | 24/12/1995 | 27/1/1996 | 34 | NW-N |
| mq1-5814-95 | particle trace |  | 113.7 | 361.1 | 8/12/1995 | 2/5/1996 | 146 | E-SE |
| mq2-20916-96 | particle trace |  | 109.4 | 248.1 | 1/12/1996 | 11/12/1996 | 10 | E-SE |
| mq2-22500-96 | particle trace |  | 199.7 | 88.6 | 18/12/1996 | 24/12/1996 | 6 | S-SW |
| mq2-26623-96 | particle trace |  | 134.4 | 413.9 | 18/12/1996 | 10/6/1997 | 174 | E-SE |
| mq2-28482-96 | particle trace |  | 98.7 | 237.4 | 19/12/1996 | 2/2/1997 | 45 | E-SE |
| mq3-17217-99 | particle trace |  | 116.0 | 378.0 | 1/1/2000 | 25/6/2000 | 176 | E-SE |
| mq3-20918-99 | particle trace |  | 139.3 | 367.9 | 11/12/1999 | 8/5/2000 | 149 | SE-S |
| mq3-22484-99 | particle trace |  | 87.8 | 864.9 | 17/12/1999 | 20/5/2000 | 155 | NE-E |
| mq3-22488-99 | particle trace |  | 80.9 | 180.8 | 16/12/1999 | 13/8/2000 | 241 | NE-E |
| mq3-22498-99 | particle trace |  | 115.2 | 313.6 | 2/12/1999 | 17/1/2000 | 46 | E-SE |
| mq3-26627-99 | particle trace |  | 133.0 | 353.8 | 14/12/1999 | 8/6/2000 | 177 | E-SE |
| mq3-26629-99 | particle trace |  | 146.6 | 483.2 | 22/12/1999 | 9/4/2000 | 109 | SE-S |
| mq3-2849-99 | particle trace |  | 131.2 | 443.0 | 27/12/1999 | 26/6/2000 | 182 | E-SE |
| mq3-28494-99 | particle trace |  | 142.5 | 454.9 | 23/12/1999 | 1/2/2000 | 40 | SE-S |
| mq3-28496-99 | particle trace |  | 112.9 | 292.2 | 24/12/1999 | 13/8/2000 | 233 | E-SE |
| mq3-28497-99 | particle trace |  | 118.8 | 269.6 | 7/12/1999 | 20/4/2000 | 135 | E-SE |
| mq3-28500-99 | particle trace |  | 93.2 | 189.9 | 17/12/1999 | 26/12/1999 | 9 | E-SE |
| mq3-28504-99 | particle trace |  | 92.9 | 278.5 | 29/12/1999 | 20/6/2000 | 174 | E-SE |
| mq4-Alice-00 | particle trace |  | 154.3 | 583.1 | 29/12/2000 | 21/7/2001 | 204 | SE-S |
| mq4-Billie-00 | particle trace |  | 130.2 | 442.4 | 28/12/2000 | 16/7/2001 | 200 | E-SE |
| mq4-Cleo-00 | particle trace |  | 146.4 | 572.4 | 30/12/2000 | 30/7/2001 | 212 | SE-S |
| mq4-Ella-00 | particle trace |  | 111.2 | 173.7 | 10/1/2001 | 12/4/2001 | 92 | E-SE |
| mq4-FirstOne-00 | particle trace |  | 136.5 | 493.0 | 24/11/2000 | 30/1/2001 | 67 | SE-S |
| mq4-Flora-00 | particle trace |  | 127.7 | 520.5 | 2/1/2001 | 26/6/2001 | 175 | E-SE |

Supplementary Table 2. Model selection out for 1^st^ trip survival using the ‘dredge’ function from MuMIn R package. Rows in bold are the candidate models used to average and get the final model because their difference in AICc is < 2.

| **model** | **(Intercept)** | **is_ESE** | **weanmass** | **year** | **is_ESE:weanmass** | **df** | **logLik** | **AICc** | **delta** | **weight** |
| --- | --- | --- | --- | --- | --- | --- | --- | --- | --- | --- |
| **2** | **-0.118** | **+** |  |  |  | **2** | **-27.2** | **58.7** | **0** | **0.328** |
| **1** | **0.56** |  |  |  |  | **1** | **-28.8** | **59.8** | **1.07** | **0.192** |
| 4 | 0.184 | + | -0.00259 |  |  | 3 | -27.2 | 60.9 | 2.22 | 0.108 |
| 6 | -42.8 | + |  | 0.0214 |  | 3 | -27.2 | 61 | 2.29 | 0.104 |
| 5 | -101 |  |  | 0.0509 |  | 2 | -28.8 | 61.9 | 3.16 | 0.0677 |
| 3 | 0.759 |  | -0.00167 |  |  | 2 | -28.8 | 61.9 | 3.23 | 0.0651 |
| 12 | -1.35 | + | 0.0105 |  | + | 4 | -26.5 | 62.1 | 3.38 | 0.0604 |
| 8 | -106 | + | -0.00396 | 0.0532 |  | 4 | -27.1 | 63.3 | 4.56 | 0.0336 |
| 7 | -163 |  | -0.00385 | 0.0823 |  | 3 | -28.7 | 64 | 5.3 | 0.0231 |
| 16 | -143 | + | 0.00906 | 0.071 | + | 5 | -26.5 | 64.5 | 5.78 | 0.0182 |

Supplementary Table 3. Model selection out for 1^st^ year survival using the ‘dredge’ function from MuMIn R package. Rows in bold are the candidate models used to average and get the final model because their difference in AICc is < 2.

| **model** | **(Intercept)** | **is_ESE** | **weanmass** | **year** | **is_ESE:weanmass** | **df** | **logLik** | **AICc** | **delta** | **weight** |
| --- | --- | --- | --- | --- | --- | --- | --- | --- | --- | --- |
| **3** | **-2.37** |  | **0.0149** |  |  | **2** | **-27.3** | **59** | **0** | **0.209** |
| **4** | **-2.96** | **+** | **0.0148** |  |  | **3** | **-26.4** | **59.4** | **0.405** | **0.17** |
| **1** | **-0.56** |  |  |  |  | **1** | **-28.8** | **59.8** | **0.812** | **0.139** |
| **2** | **-1.18** | **+** |  |  |  | **2** | **-27.8** | **59.9** | **0.973** | **0.128** |
| **12** | **-5.29** | **+** | **0.0329** |  | **+** | **4** | **-25.7** | **60.5** | **1.49** | **0.0992** |
| 7 | 41.6 |  | 0.0155 | -0.022 |  | 3 | -27.3 | 61.3 | 2.29 | 0.0663 |
| 5 | -210 |  |  | 0.105 |  | 2 | -28.6 | 61.5 | 2.51 | 0.0594 |
| 8 | 100 | + | 0.0162 | -0.0517 |  | 4 | -26.3 | 61.7 | 2.75 | 0.0528 |
| 6 | -169 | + |  | 0.084 |  | 3 | -27.7 | 61.9 | 2.97 | 0.0471 |
| 16 | 92 | + | 0.0343 | -0.0488 | + | 5 | -25.7 | 62.9 | 3.97 | 0.0287 |
